# Seeing, fast and slow: the effects of processing time on perceptual bias

**DOI:** 10.1101/556944

**Authors:** Ron Dekel, Dov Sagi

## Abstract

Fast and slow decisions exhibit distinct behavioral properties, such as the presence of decision bias in faster but not slower responses. This dichotomy is currently explained by assuming that distinct cognitive processes map to separate brain mechanisms. Here, we suggest an alternative, single-process account based on the stochastic properties of decision processes. Our experimental results show perceptual biases in a variety of tasks (specifically: learned priors, tilt illusion, and tilt aftereffect) that were much reduced with increasing reaction time. To account for this, we consider a simple yet general explanation: prior and noisy decision-related evidence are integrated serially, with evidence and noise accumulating over time (as in the standard drift diffusion model). With time, owing to noise accumulation, the prior effect is predicted to diminish. This illustrates that a clear behavioral separation – presence vs. absence of bias – may reflect a simple stochastic mechanism.

**Highlights:**

- Perceptual and decisional biases are reduced in slower decisions.
- Simple mechanistic single-process account for slow bias-free decisions.
- Signal detection theory criterion is ~zero in decision times>median.

## Introduction

A powerful idea in the neurosciences is that decision makers, brains included, integrate evidence over time to reduce error (Gold & Shadlen, 2007; Ratcliff, Smith, Brown, & McKoon, 2016). Theories adhering to this principle, such as drift diffusion models (DDM, Ratcliff, 1978), offer remarkable explanatory power, notably predicting human reaction-time on decision tasks (Ratcliff & McKoon, 2008), and accounting for neuronal activity in brain regions correlated with decision making (Gold & Shadlen, 2007). In such integrators, the initial state of accumulation is set by prior evidence favoring (biasing) one decision outcome over others (Summerfield & De Lange, 2014), implementing an approximation of Bayes’ rule (Moran, 2015). The prediction observed and investigated here is that this initial bias, strong with fast decisions, is much reduced, in some cases eliminated, with slow decisions.

### Prior-dependent bias

When faced with a difficult visual discrimination task in which one of the objects is more probable, observers tend to choose the more frequent alternative. For example, consider a task involving a fine discrimination between the orientations −0.5° and +0.5°, using briefly presented stimuli. This is a challenging task, in the sense that our observers could not always provide the correct “+” response for the “+” stimulus, showing P(answer + | +) = 0.73, and also occasionally providing the “+” response incorrectly, showing P(answer + | −) = 0.35 (mean, SEM ≤0.04; *N*=7 observers). The response bias in the task, which is the overall preference for responding “+” over “-“, can be quantified by comparing the average of these two conditional probabilities to 0.5. Here the average shows 0.54, slightly above 0.5, indicating a small bias in favor of the “+” response. (This bias may reflect a small non-significant deviation between the perceived and physical orientation in the average observer; t(6)=1.59, p=0.16.) Importantly, manipulating the proportion of +0.5° to be higher than that of −0.5° (75% of trials) leads observers to being more likely to provide the “+” response for both alternatives, which showed P(+ | +) = 0.80 and P(+ | −) = 0.44, whereas a low proportion of +0.5° (25% of trials) leads to an opposite bias, which showed P(+ | +) = 0.65 and P(+ | −) = 0.30 (mean, SEM ≤0.04; an average change from 25% to 75% of *M*=0.15 with t(6)=5.10, p=0.002, two-tailed paired t-test). Overall, these results illustrate the classic finding that decision bias is shifted in accordance with learned task priors (Gorea & Sagi, 2000; Green & Swets, 1966).

Next, we split the decision data into two equal-quantity bins, around the median reaction time (RT, reflecting the time it took the observer to provide a response; this was done separately for each experimental block and for each observer). The results showed that the overall bias measured across all trials reflects a strong bias in the faster trials (*M*=0.31, t(6)=7.91, p=0.0002, Fig. 1A) and remarkably no bias in the slower trials (*M*=-0.02, t(6)=-0.63, p=0.55, Fig. 1B). To better quantify this observation, we used a probit scale for the probability measurements, i.e., probabilities were transformed using z(⋅), the inverse cumulative distribution function of the standard normal distribution (using four equal-quantity bins, Fig. 1C). This description of behavioral data permits convenient visualization in terms of measures motivated by signal detection theory (SDT) for bias and accuracy (Green & Swets, 1966):

**Fig. 1.**
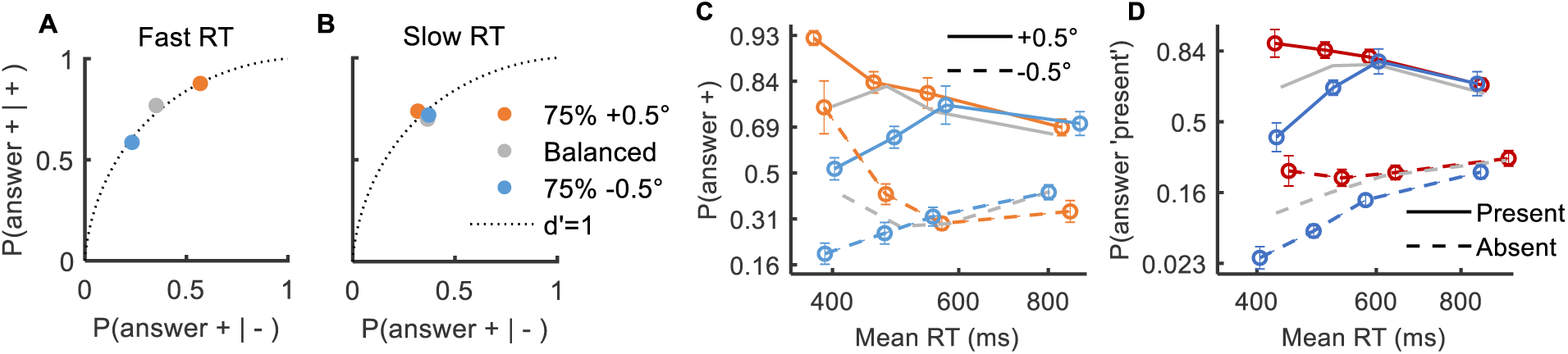
Prior-dependent bias disappears with decision time. In the experiments reported here, observers performed a visual discrimination task in which one of the objects is more probable (a prior of 25%-75%). (**A and B**) Receiver operating characteristic (ROC) plot of the probability for correct discrimination as a function of the probability for incorrect discrimination, averaged across observers, in reaction times (RTs) that are (**A**) faster or (**B**) slower than the median. (**C and D**) Response probability as a function of reaction time, for different priors (colors), and different stimuli (line styles), averaged across observers over four equal-quantity bins. (A) Discrimination experiment (as in panels A and B). (**D**) Detection experiment, target-present probability indicated by color (75%, 50%, and 25% corresponding to red, gray, and blue, respectively). The results showed a significant prior-dependent bias (a change due to color) only in fast responses. Error bars are ±1SEM.

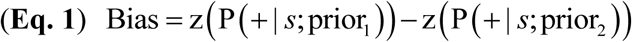

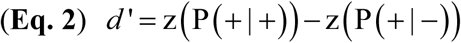

Specifically, bias (Eq. 1) is the change in response probability to a fixed stimulus *s* under different priors (the vertical distance between data points of different colors in Fig. 1C; formally equivalent to the change in internal criterion, see Supplementary materials). This measure showed that the experimental manipulation of priors led to a robust bias in fast, but not slow, trials (F(3,18)=43.66, p<0.0001, for the interaction term in a repeated-measures ANOVA of bias at the different RT bins, where bias is calculated between the 25% and 75% priors, Fig. S1AB). Similarly, accuracy (formally sensitivity or d’, Eq. 2) is the change in response probability between two stimuli under a fixed prior (the vertical distance between data points of the same color with different line styles in Fig. 1C). This measure showed that changes in accuracy were small and inconsistent (d’≈1 across priors and decision times, Fig. S1C).

Next, we consider a classic detection task, whereby observers report whether a visual target is present or absent from the display. To experimentally manipulate priors, the proportion of target-present relative to target-absent trials was changed (from 25% up to 75%). Again, the results showed a prior-dependent bias, which was strong in the fast decisions, and almost entirely absent in the slow decisions (F(3,24)=32.76, *N*=9, p<0.0001, Figs. 1D, S1DE). The accuracy showed an inconsistent but overall significant reduction with decision time (F(3,24)=7.56, p=0.001; Fig. S1F), though not by much (~40% reduction from the fastest to the slowest bin). Overall, in both discrimination and detection tasks, a significant change in behavior due to the learned prior was restricted to the faster half of the responses.

### Context-dependent bias

Motivated by the interaction of decision time and bias when manipulating the prior of the decision alternatives, we were interested in context-dependent biases, usually considered to be a perceptual effect, often considered as visual illusions. Specifically, with perhaps any visual property (orientation, luminance, color, motion, size, or facial expression, Clifford & Rhodes, 2005; Webster, 2011), the contextual value of the property leads to a bias in the reported perception of that property. For example, a physical vertical (0°) test may be reported as if it is tilted −2° due to a context of +20° surrounding the test (spatial context, Fig. 2A) or viewed before the test (temporal context; similarities of spatial and temporal contexts are discussed elsewhere, Schwartz, Hsu, & Dayan, 2007).

**Fig. 2.**
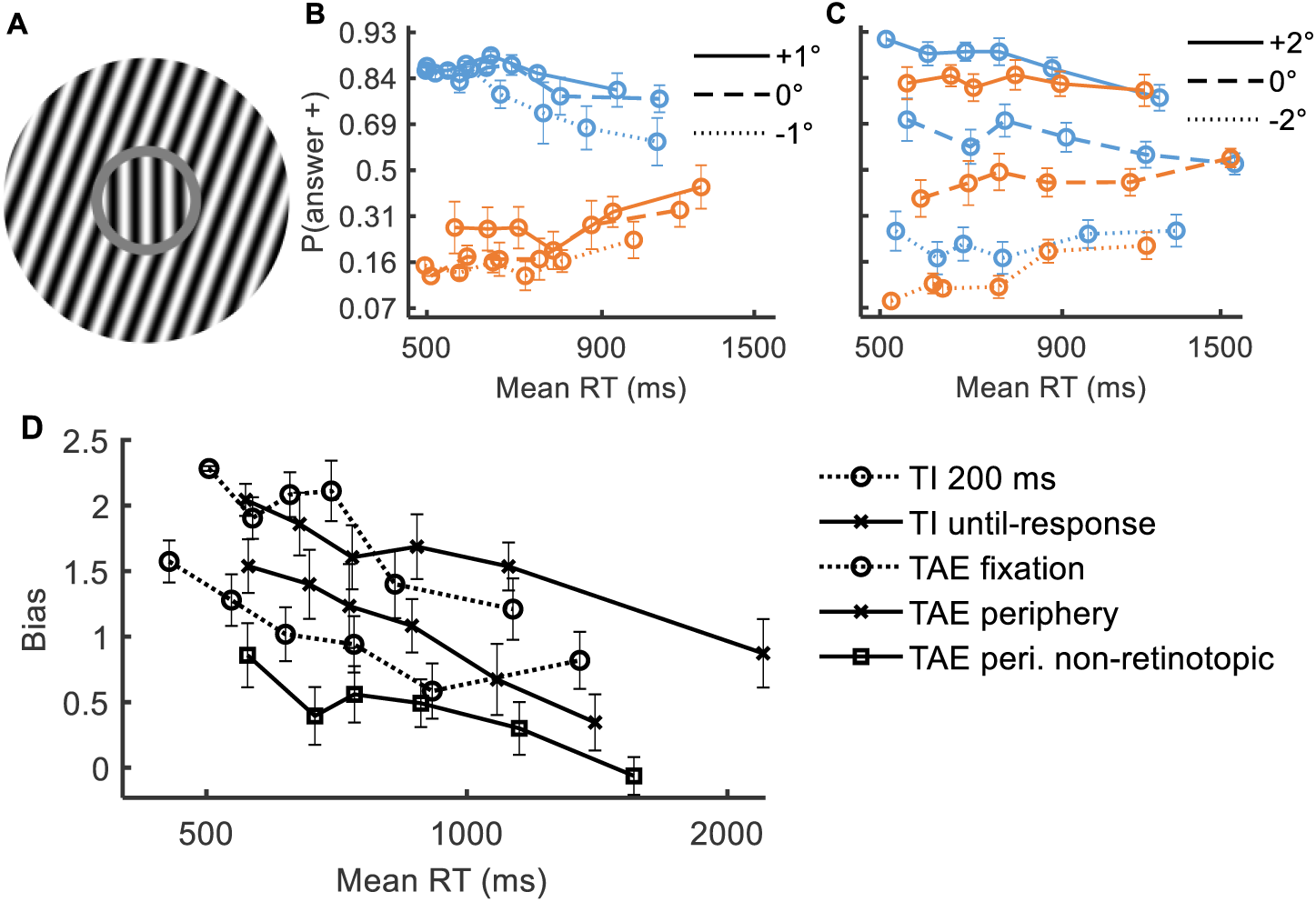
Context-dependent bias disappears with decision time. (**A**) Tilt illusion (TI, spatial context). An oriented surrounding context leads to a change in the perceived orientation of a center. (**B and C**) Response probability as a function of reaction time, under different context orientations (−20° in blue, +20° in orange), and different test orientations (line styles, see legend), averaged across observers for six approximately equal RT bins in the (**B**) TI 200 ms condition, and (**C**) tilt aftereffect (TAE, temporal context) periphery non-retinotopic condition. (D) Bias (Eq. 1) due to context orientation as a function of reaction time (six bins), for a vertical test (0°), averaged across observers. Error bars are ±1SEM.

First, we measured the influence of oriented context on a subsequently perceived orientation (tilt aftereffect, TAE). To verify that the decision time is not confounded with the presentation duration of the test (Kaneko, Anstis, & Kuriki, 2017; Wolfe, 1984), we used briefly presented stimuli (50 ms). The results showed standard effect magnitudes, with bias opposite to the orientation of context. Most importantly, the magnitude of contextual influence was reduced in slower decisions (Figs. 2C, S2). Indeed, across test orientations and conditions, the measured bias showed about 50% reduction from the fastest to the slowest bin (using 6 bins of approximately equal quantity, Figs. 2D, S3; interaction of bias for the vertical test and RT in a repeated-measures ANOVA showed fixation: F(5,60)=4.86, *N*=12, p=0.001, near-periphery: F(5,65)=6.93, *N*=14, p<0.0001, near-periphery non-retinotopic: F(5,65)=2.95, *N*=14, p=0.02; "non-retinotopic" in the sense that the adaptor and the test were presented at different retinal positions). Note that unlike the known reduction in the aftereffect magnitude with increased time difference between the adapting context and the test (Magnussen & Johnsen, 1986), here the involvement of decision mechanisms was measured using a fixed target-to-adapter time by analyzing the TAE at different reaction times.

Next, we measured the influence of oriented surrounding context on the perceived orientation of a central test (tilt illusion, TI). When the presentation of the stimulus (test+surround) persisted until the observers’ response, the results showed a clear reduction in bias for increased reaction times (bias of ~2 at ~550 ms, decreasing to bias=~0.9 at ~2000 ms; F(5,45)=3.98, *N*=10, p=0.005, Figs. 2D, S3). Importantly, when using a fixed presentation duration (200 ms, briefer than any decision), the results again showed a reduction in bias with time (bias of ~2.3 at ~500 ms, decreasing to bias=~1.2 at ~1000 ms; F(5,45)=8.71, *N*=10, p<0.0001, Figs. 2BD, S3). The results indicate that the observed reduction in illusion magnitude is mediated by changes in the decision-making processes.

The accuracy in the tasks showed an inconsistent and mostly minor reduction in slower decisions (as measured by reduced d’, Fig. S3), suggesting a dissociation between change in bias and change in accuracy at different reaction times (moreover, reductions in accuracy are typically associated with increased rather than decreased bias, Wei & Stocker, 2017). An alternative measure for bias, though less accurate (see the Supplementary materials) is the shift in the perceived vertical orientation measured in degrees. Results using this measure showed similar results as reported above (Fig. S4). Overall, the magnitudes of both TAE and TI were reduced in slower decisions, revealing an interaction between contextual influence and decision-making processes.

### Theory

Next, we aimed to account for the observed reduction in bias with decision time by applying general principles. Generally, a system that accumulates noisy evidence when making a decision can be interpreted as a stochastic decision process (for example, Dayan & Daw, 2008). Interestingly, in many stochastic processes (notably “memoryless” ones, such as random walks, Markov chains, and typical diffusion models), with processing time, the process gradually becomes independent of the initial state due to accumulated stochasticity (noise), so biases that reflect initial conditions are expected to gradually decrease with time. For example, in simple unbounded diffusion (random walk), the initial state is lost at a slow rate proportional to 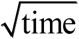 (Fig. 3A), as in many stochastic processes. From the Bayesian perspective, the initial state measures the a priori information, and this prior is outweighed when more noisy evidence has been accumulated, leading to reduced prior effect in slower decisions.

**Fig. 3.**
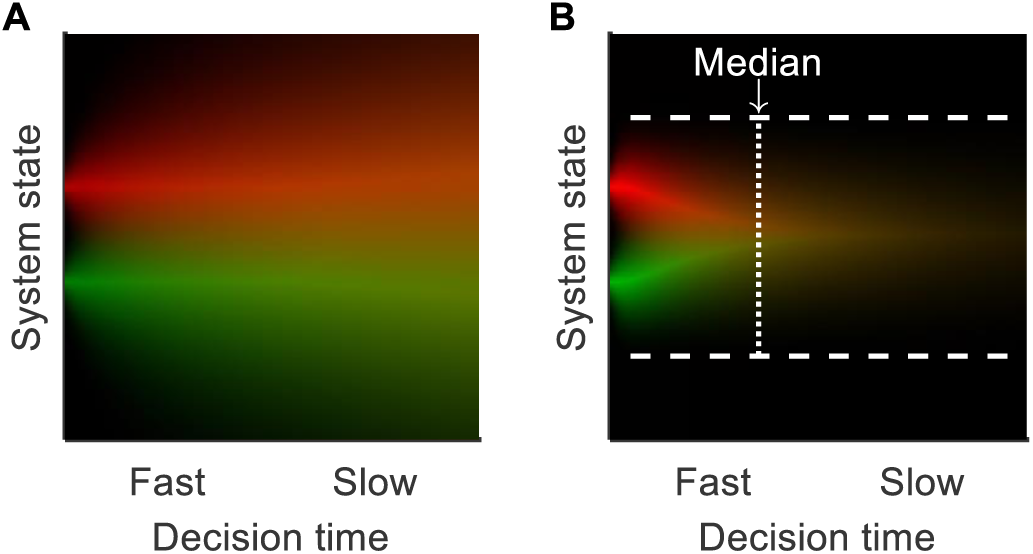
Bias reduction due to accumulated noise. Probability density function (pdf) of the system state at different times, for different initial conditions (i.e., priors; red vs. green), in (**A**) a diffusion process with no bounds, and in (**B**) a diffusion process that stops when a bound (dashed lines) is reached. The effect of the initial state is lost with time (yellow color mix).

Importantly, decision bias is reduced rapidly (~exponentially) in bounded decision models such as the Drift Diffusion Model (DDM, Ratcliff, 1978), in which the process of evidence accumulation continues until a bound is reached (Fig. 3B). For a starting point that is not too extreme (i.e., a moderate bias), the model predicts almost zero bias after the median decision time (Fig. 4). Indeed, the model provides an excellent account (Figs. 4, S5) for the rapid reduction of prior-dependent bias observed here (Fig. 1). This result is consistent with earlier modeling of experimental data with the DDM (Mulder, Wagenmakers, Ratcliff, Boekel, & Forstmann, 2012; Summerfield & De Lange, 2014; White & Poldrack, 2014). Note that the theory presented here to explain decision time-dependent bias can also account for experimental situations where the bias effect persists with time (e.g., a change in the drift rate in the DDM), as evident with experimental manipulations affecting external noise selection (Urai, De Gee, Tsetsos, & Donner, 2018). More generally, persistent evidence selection should clearly lead to persistent bias effects (Kloosterman et al., 2019; White & Poldrack, 2014).

**Fig. 4.**
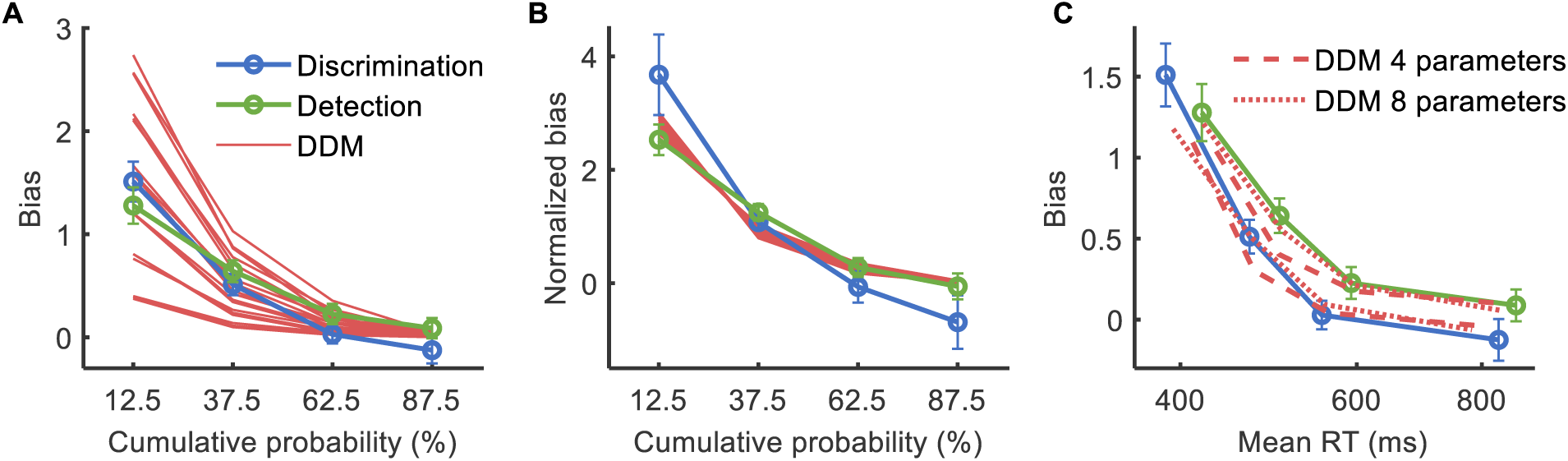
Reduction in prior-dependent bias is explained by the drift diffusion model. (**A**) Bias (Eq. 1) as a function of cumulative probability, for behavioral data of the prior-dependent experiment (averages of Fig. S1AB), and for a change in starting point in the DDM measured in four RT bins (red lines; the different groups of superimposed lines reflect different sizes of starting point changes away from the mid-point, and superimposed lines reflect varying drift rates; the influence of the bound-separation parameter is ignored because, for reasonable parameter values, it is mostly accounted for by changes in drift rate and starting point, by virtue of considering cumulative probability). (**B**) Bias from panel A divided by the average value (calculated separately for each observer and then averaged across observes). It can be seen that the stereotypical rate of reduction of bias owing to a starting point change in the DDM is qualitatively similar to behavior. (**C**) Bias as a function of RT, in behavior, and for DDM fitted to behavior based on the RT distributions (Fig. S5), using 4 or 8 fitting parameters (see Methods), separately for each observer, then averaged across observers. Error bars are ±1SEM.

With context-dependent bias, the observed reduction in bias was slow (Fig. 2D). This can be explained as a partially persistent bias effect (e.g., a context-dependent change in both prior and the rate of evidence accumulation, which is addressed within the DDM framework, Figs. 5, S6), or as an unbounded diffusion process (i.e., the 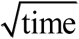 rate of bias decay, consistent with decision timing that is independent of the system state, Fig. 5). Note that the sign of the context-dependent bias is opposite to the prior (“anti-Bayesian”), consistent with a shift of the reference (i.e., bound positions) in the direction of the prior. A seemingly promising alternative to the above interpretations is that the influence of adapting context is time independent but varies across trials. However, this would predict the absence of line intersections in plots of reported probability vs. time (Fig. S7), unlike behavioral data (Figs. 1B, S2). More generally, preferring one mechanistic model over another is within the boundaries of the explanation we offer.

**Fig. 5.**
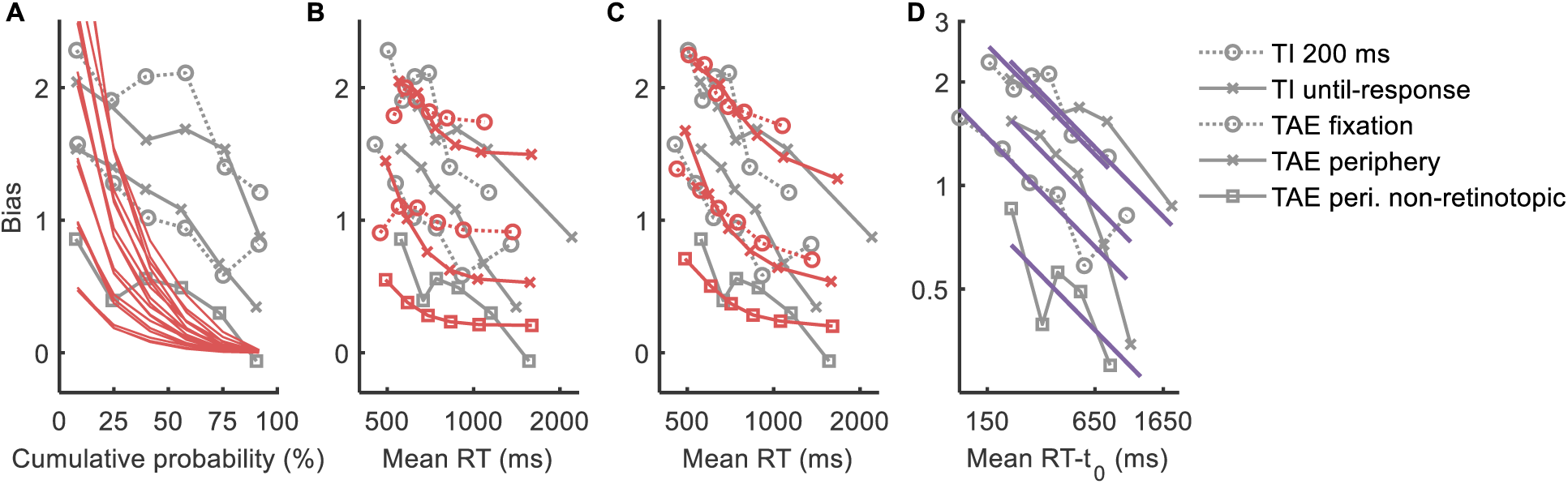
Modeling the reduction in context-dependent bias. Shown is bias (Eq. 1) in models and in the behavioral context experiments (gray lines, reproduced from Fig. 2D). (**A-C**) DDM model, measured in six RT bins (red lines). (**A**) Bias as a function of cumulative RT probability, where modeled DDM bias is caused by a change in starting point (for different fixed values of drift rate and for different sizes of starting point change, as in Fig. 4A). The behavioral rate of reduction in bias is much slower than modeled. (**B**) Bias as a function of RT, where modeled bias is the DDM fit to the behavioral RT distribution, averaged across observers (see Fig. S6), using 4 fitted parameters. (**C**) Same, using 8 fitted parameters. (**D**) Bias as a function of *RT* – *t*_0_ in a log-log plot, in behavior, and the least-squares fit to a 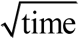 decay rate (purple lines; possibly reflecting a fixed rate of noise accumulation as in an unbounded random walk). The *t*_0_ measures the non-decision time, presumably mostly post-decision, which was set to a fixed value of 350 ms. Overall, the different modeling approaches can qualitatively account for a slow rate of reduction in bias. Error bars are ±1SEM.

## Discussion

Overall, this work illustrates a novel way of thinking about bias and time in human decision-making. Instead of fitting a specific model to data (as in White & Poldrack, 2014), we make a simple yet powerful general claim: that bias derived from starting conditions (e.g., prior) gradually decreases with decision time. This claim applies to an entire family of stochastic decision processes, which can be indistinguishable with limited data, emphasizing the importance of focusing on the general principle.

Most importantly, the finding of reduced bias with longer decision times may appear to perfectly conform with a dual-theory account of a transition between separate systems (Evans & Stanovich, 2013; Kahneman, 2011; Tversky & Kahneman, 1974): from a fast system that is bias-prone, to a slow system that is bias-free. Similarly, the known reduction in context-dependent bias with the duration of testing stimulus presentations can currently be explained by assuming dual processing (in low vs. high systems) (Kaneko et al., 2017; Wolfe, 1984). However, we have found that using models based on the decision principles described above offers a full mechanistic explanation. Even a rapid reduction in bias (with the slower half of responses measuring practically zero bias) can be explained by a rule for when to stop accumulating evidence that depends on the extent of evidence that has been accumulated, an account with very strong support in brain decision making (DDM, Gold & Shadlen, 2007; Ratcliff & McKoon, 2008; Ratcliff et al., 2016) and clear statistical implications (SPRT, Moran, 2015; Summerfield & De Lange, 2014). Of course, there can be dual processes in the brain, and there are low vs. high brain areas, but at least for the basic perceptual phenomena considered here, there is a simple mechanistic account that does not need to assume multiple systems.

Traditionally, perceptual phenomena are considered to be a part of the faster system (Tversky & Kahneman, 1974), although there is no clear definition. One attempt at formulating a definition by Evans and Stanovich (Evans & Stanovich, 2013) suggests that “rapid autonomous processes (Type 1) are assumed to yield default responses unless intervened on by distinctive higher order reasoning processes (Type 2)”, which at least phenomenologically appear to match the reduction in bias discussed here. Regardless of terminology, perceptual decisions and the simple mechanistic models they permit should be considered in the discussion about fast vs. slow systems.

## Materials and Methods

The purpose of this study was to investigate the interaction between bias and decision time for perceptual and decisional biases.

### Observers

Thirty-three observers (25 females, 8 males, aged 18-31) with normal or corrected-to-normal vision participated in the experiments. All observers were naïve to the purpose of the experiments, and provided their written informed consent. The work was carried out in accordance with the Code of Ethics of the World Medical Association (Declaration of Helsinki). Most observers had prior experience in participating in psychophysical experiments.

### Apparatus

The stimuli were presented on a 22” HP p1230 monitor operating at 85Hz with a resolution of 1600×1200 that was gamma-corrected (linearized). The mean luminance of the display was 26.06 cd/m^2^ (detection, discrimination, and TAE experiments) or 48.96 cd/m^2^ (TI experiments) in an otherwise dark environment. The monitor was viewed at a distance of 100 cm.

### Stimuli and tasks

All stimuli were presented using dedicated software on a uniform gray background. To begin stimulus presentation in a trial, observers fixated on the center of the display and pressed the spacebar. Responses were provided using the left and right arrow keys. Distances are reported in degrees (°) of the visual field. As described below, the used Gabor patches were of the same parametrization (except orientation, monitor position, and contrast).

#### Discrimination (2AFC) experiment

Stimuli were Gabor patch targets tilted −0.5° or +0.5° relative to vertical (50 ms presentation, 50% contrast, σ=0.42°, λ=0.3°, random phase, 750 ms onset ± up to 100 ms onset jitter). Observers were instructed to report whether the orientation of the target is clockwise or counter-clockwise to vertical (2AFC), with auditory feedback indicating mistaken reports. Four peripheral crosses co-appeared with the target.

#### Detection experiment

In the target-present trials, the stimuli were low-contrast Gabor patches (50 ms presentation, 0.5% to 1% Michelson contrast (i.e., Gabor amplitude of 0.5 to 1.25), vertical orientation, σ=0.42°, λ=0.3°, random phase, 750 ms onset ± up to 100 ms onset jitter). In the target-absent trials, nothing was presented at the center of the display. Observers were instructed to report whether the target appeared or not, with auditory feedback indicating mistaken reports. Four peripheral crosses co-appeared with the target presentation interval (in both target-present and target-absent trials).

#### TAE experiments

The following presentation sequence was used: a blank screen (600 ms presentation), Gabor “adaptor” (i.e. context, oriented −20° or +20° to vertical, 50 ms), a blank screen (600 ms), and a near-vertical Gabor “test” (50 ms). Observers were instructed to inspect the adaptor and target presentations, and then to report whether the orientation of the target was clockwise or counter-clockwise to vertical (2AFC, no feedback). Gabor patches were 50% Michelson contrast with σ=0.42°, λ=0.3°, and random phase. Two versions of the experiment were run: fixation and periphery. In the fixation experiment, adaptors and tests were presented at the fixated center of the display, and tests were oriented −9° to +9° (in steps of 1°). In the periphery experiment, adaptors and tests were presented at either left or right of the fixation (at ±1.8°, i.e. 12λ separation). The test was presented either at the same side as the adaptor (retinotopic) or at the opposite side (non-retinotopic), randomly. Tests were oriented from −12° to +12° (in steps of 2°). The reason that the TAE experiments had no variability in the onset of the test is that the time difference between adaptor and test is known to affect the magnitude of the TAE (Greenlee, Georgeson, Magnussen, & Harris, 1991). Four peripheral crosses co-appeared with the target to facilitate discrimination between the adaptor and the test.

#### TI experiments

Stimuli (e.g., Fig. 2A) consisted of a near-vertical sine-wave circular “test” (oriented from −9° to +9° (in steps of 1°), λ=0.3°, random phase, a radius of 0.6°), and a sine-wave “surround” ring (oriented −20° or +20°, λ=0.3°, random phase, width of 1.2°, and a gap of 0.15° from the central circle). Observers were instructed to inspect the circle presentation, and to report whether the orientation of the circle is clockwise or counter-clockwise to vertical (2AFC, no feedback). The test+surround stimuli were presented starting from 450 ms after the trial initiation (± up to 100 ms jitter), for a duration of either 200 ms (“200 ms” experiment) or for however long it took the observer to provide a response (“until-response” experiment).

### Procedure

In all experiments, we required a very low rate of finger errors (less than one reported mistake in 200 trials), to minimize contamination by fast guesses (Ratcliff & Tuerlinckx, 2002). All observers managed to achieve this level of accuracy with persistent coaxing. Observers were encouraged to reply quickly, unless the speed of reply led to mistakes. Most observers participated in multiple experiments of this study, separated by at least three days of a break. Under all conditions, each daily session was preceded by a brief practice block with easy stimuli (this practice was repeated until close-to-perfect accuracy was achieved).

#### Discrimination (2AFC) experiment

Observers (*N*=7) completed a single daily session. Trials were blocked into streams of 80 trials with a given prior (either 75% vs. 25%, 50% vs. 50%, or 25% vs. 75% for the −0.5° vs. +0.5° orientation alternatives). In each daily session, a set of six blocks (two per prior, in random order, each lasting ~2 minutes, separated by a 20-second break to minimize inter-block contamination by learned priors) was completed twice (totaling four blocks per prior; ~13.5 minutes per set, separated by a 2-minute break). Observers were informed that different blocks may have different priors of the decision alternatives. Two observers were disqualified before data analysis, one having anomalously high accuracy (d’=2.5, with other observers showing d’ =1.01±0.18, mean±STD), and one having an anomalously strong baseline response bias in favor of one of the alternatives, which saturates the measured probabilities. (Both disqualified observers exhibited a prior-dependent bias in fast replies and no prior-dependent bias in slow replies.) Note that the experimental design assumes that all observers have comparable sensitivity. The horizontal orientation of the monitor and the table was verified using a spirit level.

#### Detection experiment

Observers (*N*=9) completed from two to three daily sessions. Trials were blocked into streams of 80 trials with a given prior (either 75% vs. 25%, 50% vs. 50%, or 25% vs. 75% for target-present vs. target-absent). In each daily session, two sets of six blocks were completed (two per prior, in random order). Blocks (lasting ~2 minutes) were separated by a 15-second break. The two sets (~14.5 minutes each) were separated by a 2-minute break. To ensure a comparable d’ between observers and daily repetitions, the level of difficulty was adjusted per observer at the beginning of each daily session with a staircase procedure, and occasionally also during the break between the two sets.

#### TAE experiments

Sessions were composed of blocks of 125 trials (lasting ~5 minutes), separated by 2-minute breaks of blank screen free viewing. In the fixation experiment, observers (*N*=12) performed a single daily session consisting of six blocks. In the periphery experiment, observers (*N*=14) performed 3-8 daily sessions each consisting of five blocks.

#### TI experiments

Sessions consisted of blocks of 190 trials (lasting ~5 minutes), separated by 2-minute breaks of blank screen free viewing. Observers (*N*=10 and *N*=10, in “200 ms” and “until-response” experiments, respectively) performed a single daily session consisting of five blocks.

### Analysis

#### Data binning

To measure the interaction of time and bias, behavioral data were binned, based on reaction times (RT) into N bins. For the detection and discrimination experiments, binning was carried out separately for each combination of experimental block × trial type (i.e., separately for each decision alternative; note that different blocks correspond to different experimental priors). For the TAE and TI experiments, binning was carried out separately for each combination of experimental day × test orientation × context orientation (adaptor or surround). Under the TAE periphery conditions (retinotopic and non-retinotopic), binning combinations were further conditioned based on adaptor side × test side. For example, if a given combination has two trials with a RT of 350 and 500 ms, then to achieve binning for *N*=2 bins, the trial with 350 ms is put into the faster bin, and the trial with 500 ms is put into the slower bin. When the number of trials is not an exact multiple of the required number of bins, a deterministic rule was used. Binned trials were pooled from all relevant repetitions, separately for each observer. Conceptually, the strict binning rule we applied prevents interaction between trial type and the bin used (because different trial types can have different inherent difficulties and hence, different average reaction times). In practice, any reasonable binning rule we tried that had more trials than binning combinations led to nearly identical results. In the detection and discrimination experiments, the first ten trials were excluded from the analysis, so that the priors could be learned (which takes at least a few trials to learn, e.g. Zylberberg, Wolpert, & Shadlen, 2018). In the context experiments, trials with RT<300 ms were pruned (this rule addressed the rare occasions where the observers judged the perceived orientation of the adaptor instead of the test).

#### Bias and accuracy calculation

To quantify bias and accuracy, we relied on measures motivated by signal detection theory (SDT, Green & Swets, 1966). Specifically, the definition of bias (Eq. 1) is equivalent to the change in the internal criterion ^*c*^ (Eq. S1, from SDT) between two conditions.

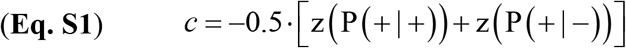

Formally, define 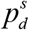 to be the response probability (the percent of ‘+’ responses) for stimulus *s* (*s* = ‘−’ or *s* = ‘+’) in condition *q* (*q* = 1 or *q* = 2, for two prior probabilities or two context orientations). Then it holds (Eq. S2) that the average of the bias of the two stimuli (i.e., 0.5 ⋅(Bias^+^ + Bias^−^) where Bias^*s*^ is the bias with stimulus *s*), is the change in the internal criterion of the two conditions (i.e., *c*_2_ – *c*_1_ where *c_d_* is the internal criterion in condition *q*).

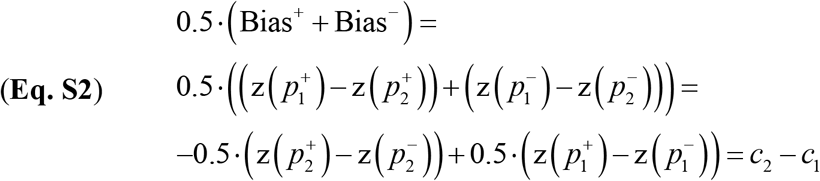

In addition, to avoid saturation, probabilities were clipped to the range 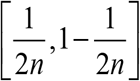, where *n* is the number of trials in the measurement. Accuracy (d’) was calculated by applying Eq. 2 after the probabilities were clipped to the above range. (An alternative approach for avoiding saturation is to average probabilities across observers before calculating bias and accuracy. This approach led to the same qualitative findings as reported here, in all experiments.)

#### Fitting perceived orientation

In addition to the bias measure motivated by SDT (described above), we also considered the magnitude of the shift in the perceived test orientation to quantify context-dependent bias. Specifically, the magnitude was defined as half the shift in the perceived vertical orientation between the two adaptor orientations. To find the orientation that is perceived as vertical in a given condition, the percentage of clockwise reports as a function of the target orientation was interpolated to find the orientation with 50% clockwise reports (i.e., equal probability for clockwise and counter-clockwise reports; fitting to a cumulative normal distribution that takes into account the lapse rates was achieved with the Psignifit 3.0 software, Fründ, Haenel, & Wichmann, 2011). Under the TAE periphery conditions (retinotopic and non-retinotopic), the data reflected two ‘test’ sides measured at several experimental days. The effect magnitude was calculated separately for the different experimental days and sides, then averaged across days and sides. Note that although this method of analysis is standard for estimating TAE and TI effect sizes, it is less meaningful when binning reaction times, because different test orientations have different difficulties, and hence, different mean decision times. Specifically, if binning is done based on time irrespective of test orientation, then the bins are unbalanced (e.g., the fastest bin will only contain trials of easy test orientations). If binning is done separately for each test orientation (as we do here), then the bins are balanced, but there is no descriptive time range associated with a bin (i.e., bins reflect relative rather than absolute times).

#### Drift diffusion model analysis

To obtain analytical function computation and fitting of the drift diffusion model (DDM, Ratcliff, 1978), we used the Fast-DM software (Voss & Voss, 2007). Fitting was achieved with a Kolmogorov-Smirnov (KS) setting, which finds the set of parameter values that minimizes the KS distance between the modeled and the behavioral cumulative RT distributions. The following model parameters were considered: starting point (normalized to the bounds separation, i.e. taking values in the range of 0 to 1), drift rate, non-decision time (in ms), bounds separation, and the asymmetry in non-decision time between the upper vs. the lower bound. Parameters reflecting the inter-trial variability of starting point, drift rate, and non-decision time were also considered (measured in units of standard deviation) (Ratcliff & Rouder, 1998). The fitting of the drift rate and of the variability in drift rate parameters permitted different fitted values for different experimental stimuli, unlike the remaining parameters that were assumed to be stimuli-independent (but still dependent on prior or context). In all reported conditions, the DDM provided a reasonably good fit of the behavioral RT distribution (typical lack-of-fit p-values of about 0.5). For the context experiments, only three test orientations were used in the fit (-k°, 0°, and +k°; k=1 for TI and TAE fixation, whereas k=2 for TAE periphery). To deal with RT contaminants, the observers were instructed to avoid finger errors (see above), and trials with extreme RT values were pruned before fitting (Ratcliff & Tuerlinckx, 2002). (The reported findings remained the same if none of the trials were pruned.) The illustrations in Fig. 3ab were obtained by simulations (100,000 trials).

---

**Fig. S1.**
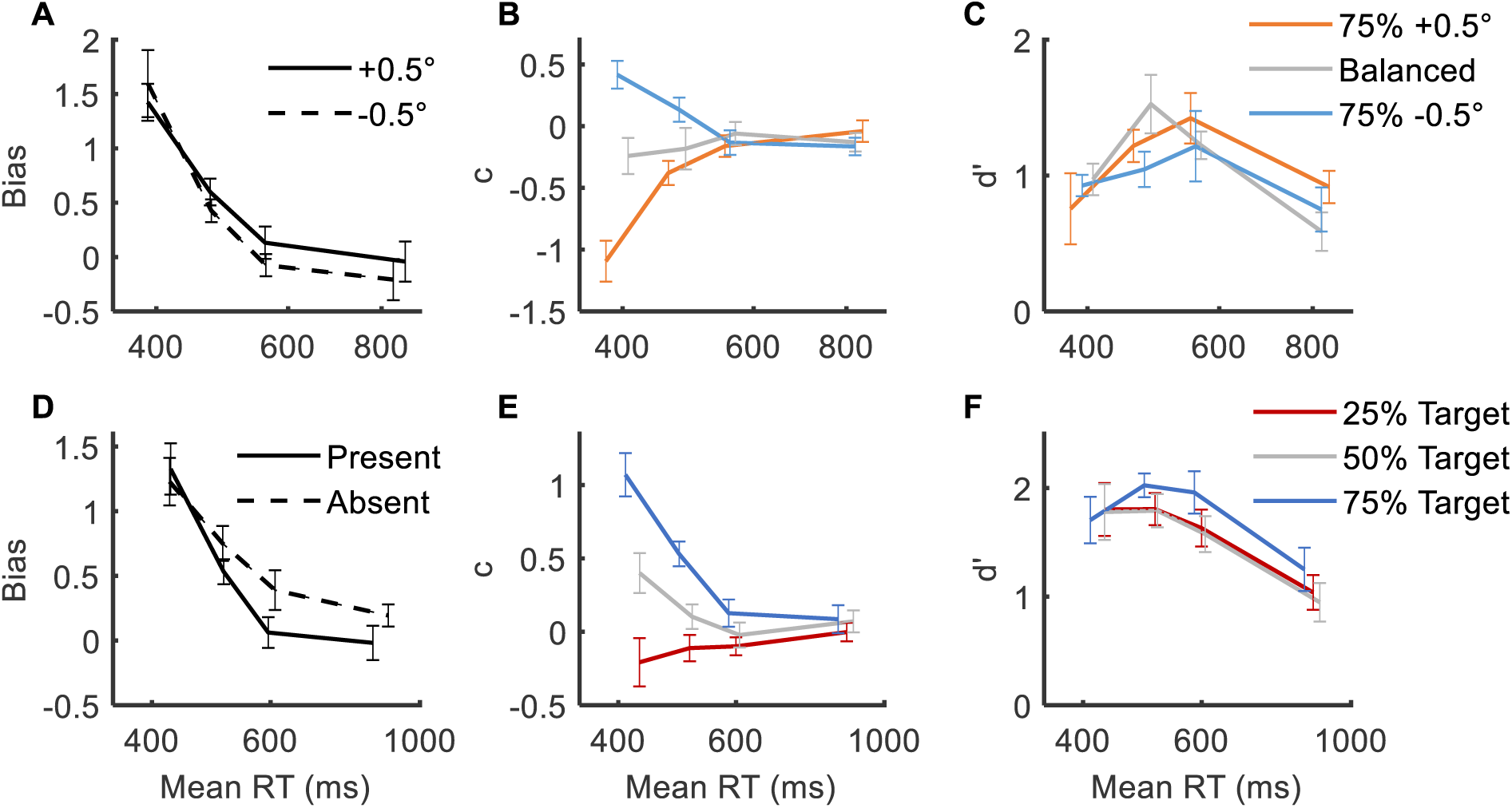
Bias, criterion, and sensitivity in the discrimination and detection tasks. (**A-C**) Shown for the discrimination task are average across observers of (**A**) bias (Eq. 1), (**B**) internal criterion (c, Eq. S1), and (**C**) sensitivity (d’, Eq. 2). (**D-F**) Same for the detection task. Note that in the detection task, the criterion data showed an asymmetry between the 25% and the 75% priors, possibly reflecting an asymmetry in learned prior: missed targets may be more likely to be interpreted as absent, despite an auditory feedback (i.e., priors are learned with a shift toward target-absent, e.g., the 50% prior is learned as if it is 40%). Error bars are ±1SEM.

**Fig. S2.**
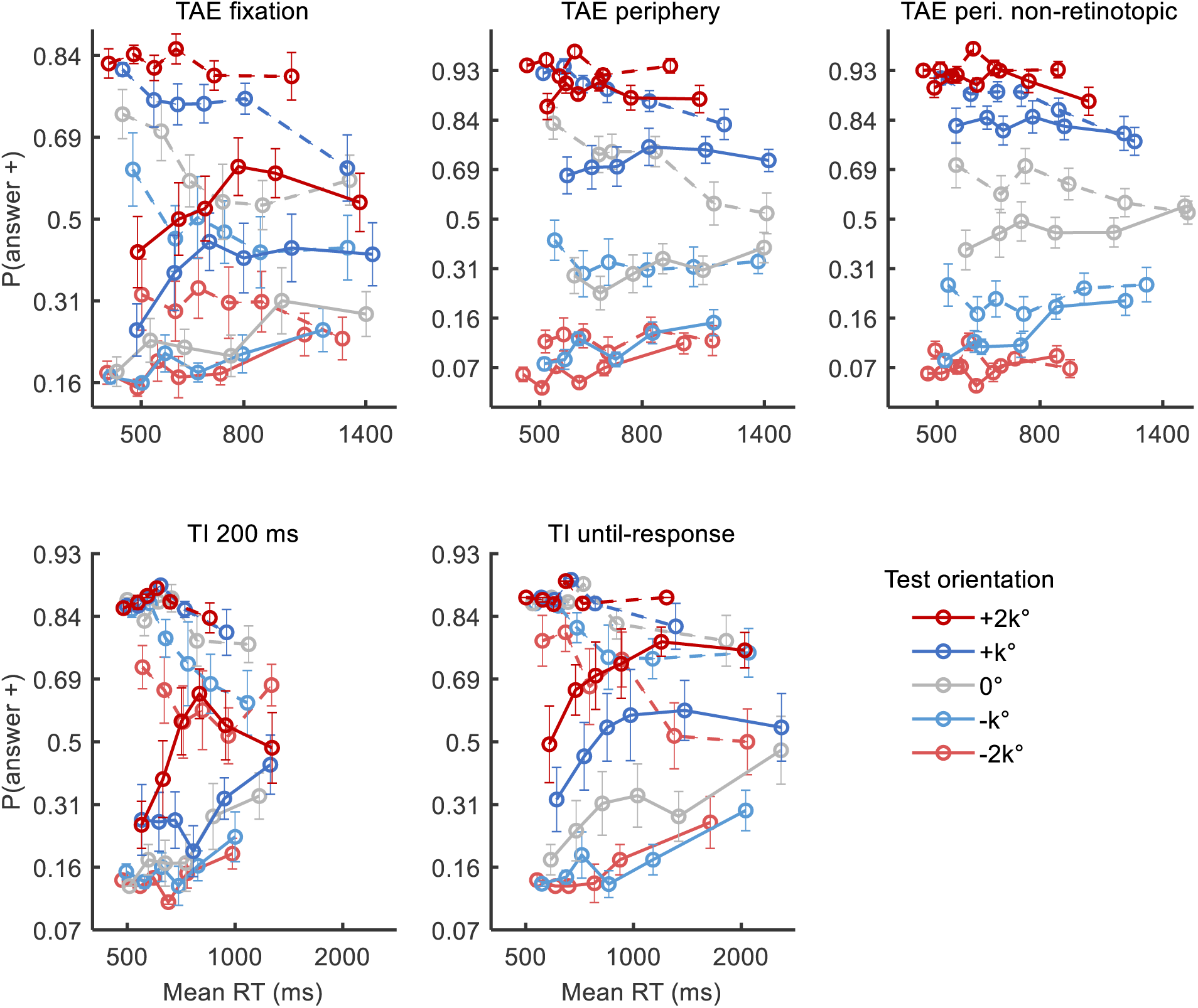
Interaction of bias and time for TAE and TI. Shown is the response probability as a function of reaction time, under different context orientations (line styles: −20° is dashed, +20° is solid), and different test orientations (colors: gray, blue, and red indicate 0°, ±k°, and ±2k° test orientations; k=1 for TI and TAE fixation, and k=2 for TAE periphery). The data in the TAE non-retinotopic and TI 200 ms conditions are partially reproduced from Fig. 2BC (with different styles). The presence of bias is denoted by a difference between the solid and dashed lines of equal color. The presence of intersections between lines corresponding to different test (color) and context (line style) orientations suggests that the context-dependent bias was time-dependent (e.g., an intersection between the dashed gray line and solid blue line in the TAE periphery condition can be taken as evidence that the 0° test with a clockwise context-induced bias is not interchangeable with the 2° test with a counter-clockwise bias). Error bars are ±1SEM.

**Fig. S3.**
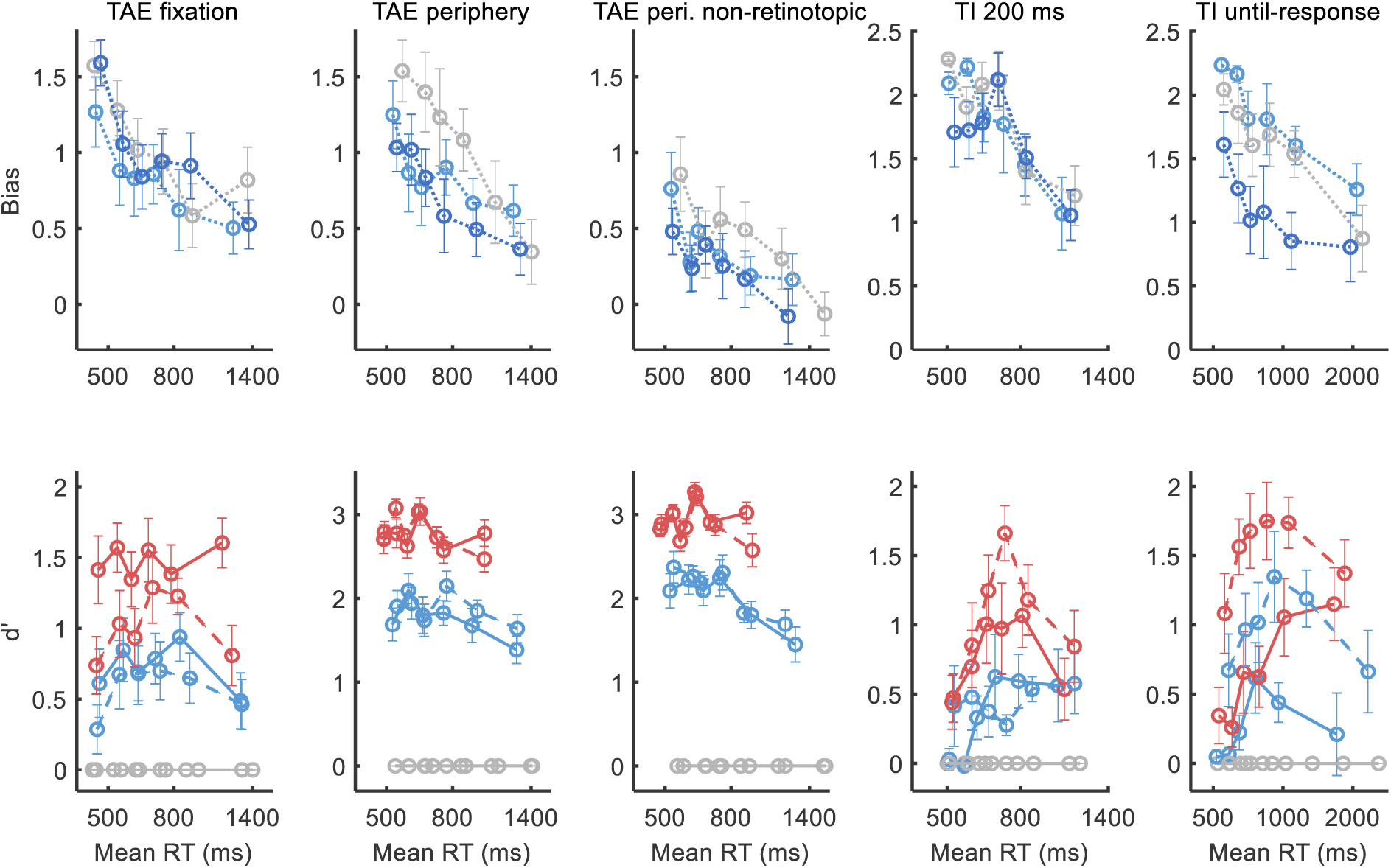
Bias and sensitivity for TAE and TI. Shown is the average across observers of bias (Eq. 1) and sensitivity (d’, Eq. 2) under the different context conditions. The different test orientations were denoted by color (gray, blue, and red indicate 0°, ±k°, and ±2k° test orientations; k=1 for TI and TAE at fixation versions, whereas k=2 for retinotopic and non-retinotopic peripheral TAE versions). For d’, measured for the +k° vs. −k° test orientation, the different context orientations were denoted by line style (−20° is dashed, +20° is solid). Results show reduction in bias across conditions and reaction times. Error bars are ±1SEM.

**Fig. S4.**
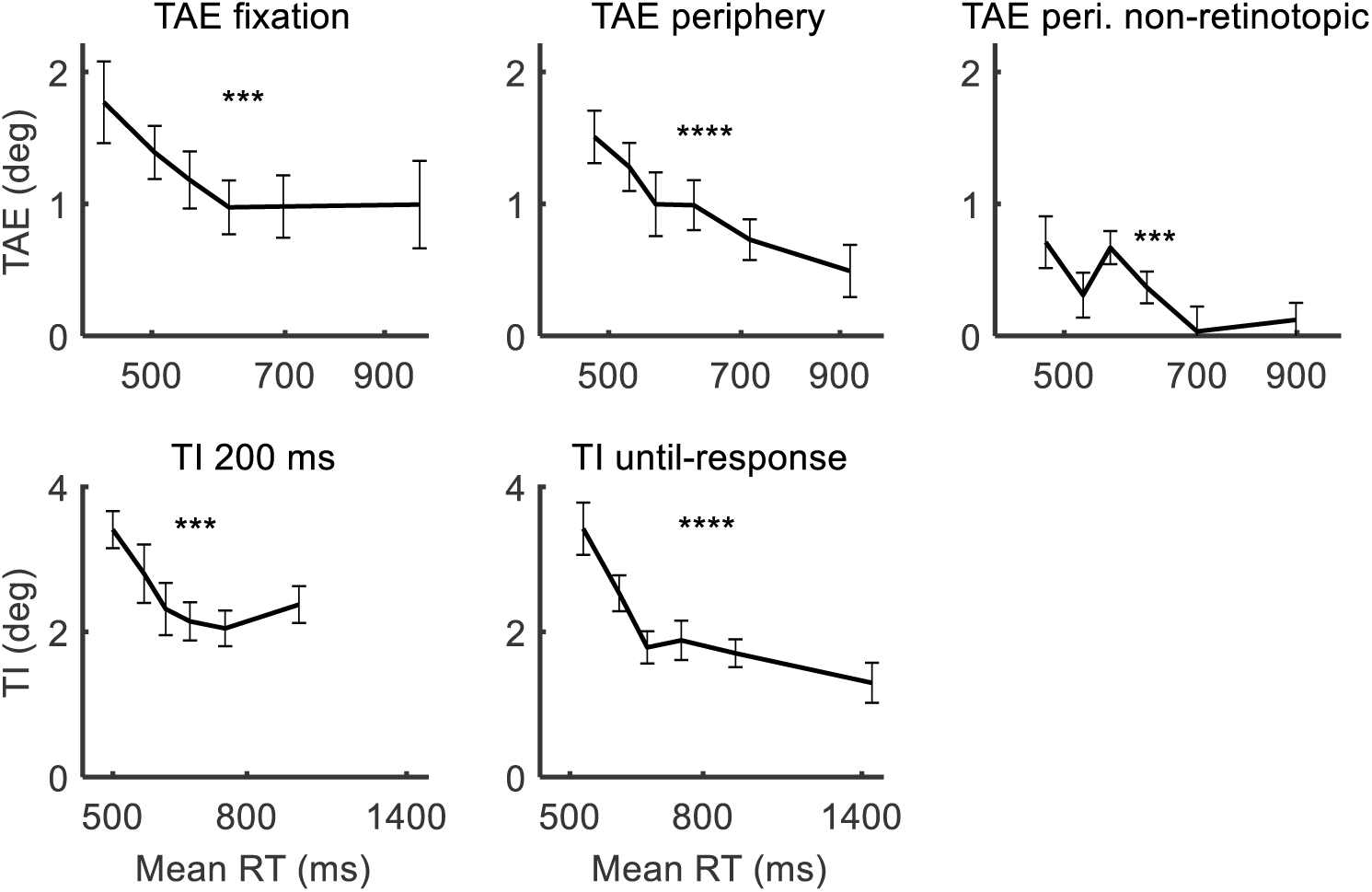
Magnitude of TAE and TI in degrees. Shown is half of the interpolated shift in the perceived vertical test orientation due to context orientation (adaptor or surround). Measurements were obtained by fitting a cumulative normal distribution to the psychometric function of percent clockwise reports as a function of test orientation (see Methods). Note that this method of analysis is less accurate than Eq. 1 for analyzing an interaction with RT, because different test orientations reflect different difficulty levels, hence different RTs. Still, to ensure a balanced number of trials per test orientation in an RT bin, the binning here was performed separately for each combination of context orientation and test orientation. Error bars are ±1SEM, and asterisks indicate the significance level of the change in magnitude in different RTs obtained with a repeated-measures ANOVA (*** p<0.001, **** p<0.0001).

**Fig. S5.**
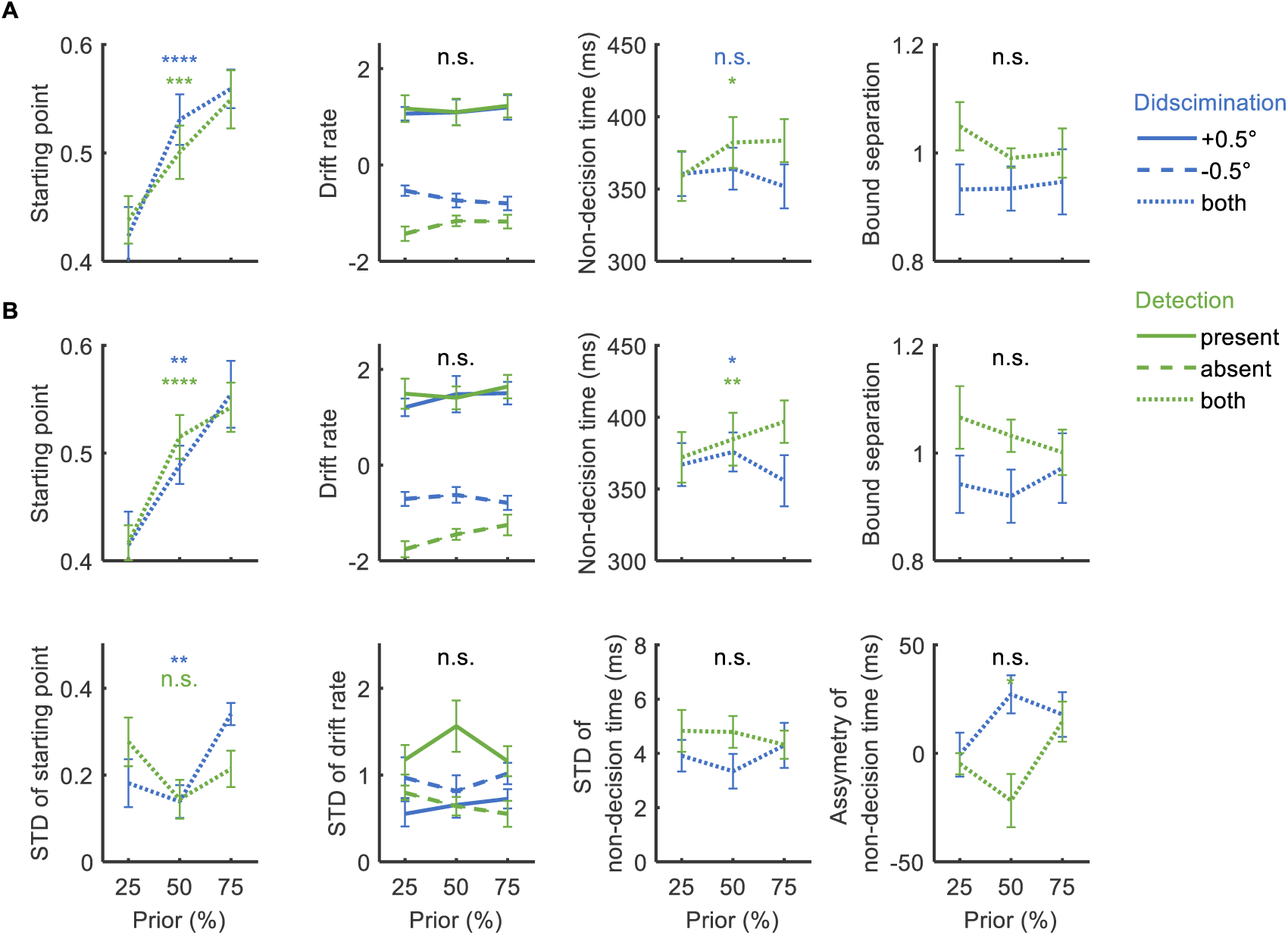
Fitted drift diffusion model parameter values in the prior experiments. Shown is the average across observers of the parameter values fitted to the behavioral RT distributions, for the different experimental priors, in the discrimination experiment (blue, the prior is for +), and in the detection experiment (green, the prior is for target). The fitted DDM had either (**A**) 4 parameters, or (**B**) 8 parameters (see Methods). Asterisks indicate the significance level of a change in parameter value with experimental prior in a repeated-measures ANOVA (two-way in the case of the drift rate and the STD of drift rate parameters). It can be seen that a change in prior was mainly explained in the DDM as a change in starting point. Annotations: n.s. not significant, * p<0.05, ** p<0.01, *** p<0.001, **** p<0.0001. Error bars are ±1SEM.

**Fig. S6.**
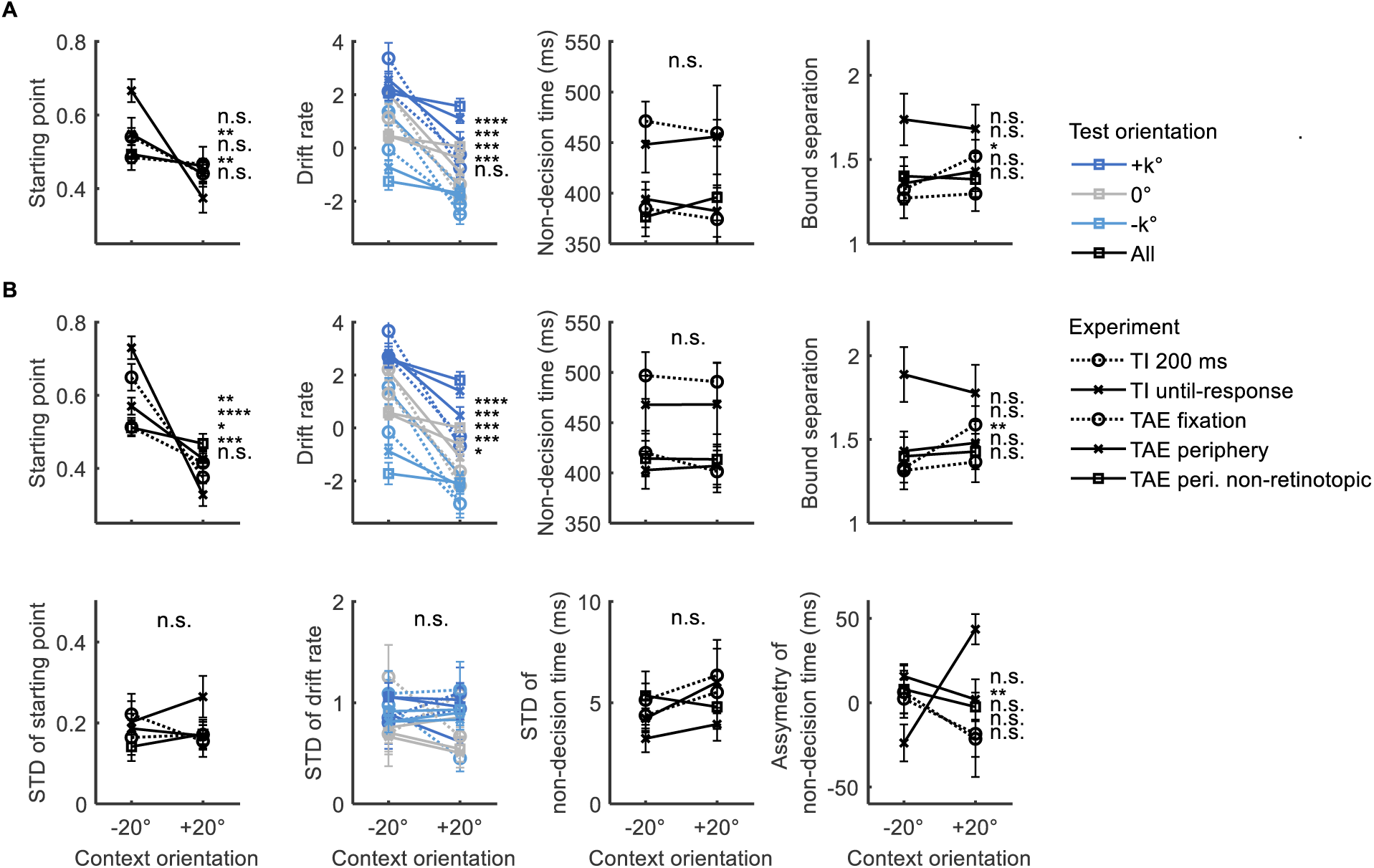
Fitted drift diffusion model parameter values in the context experiments. Shown is the average across observers of the parameter values fitted to the behavioral RT distributions for a DDM with (**A**) 4 parameters, or (**B**) 8 parameters. Only the three test orientations shown were used in the fit (k=1 for TI and TAE fixation, and k=2 for TAE periphery). Asterisks indicate the significance level of a change in parameter value with context (as in Fig. S5; vertical order following legend). It can be seen that a change in context orientation was explained in the DDM as a change of both starting point and drift rate. Error bars are ±1SEM.

**Fig. S7.**
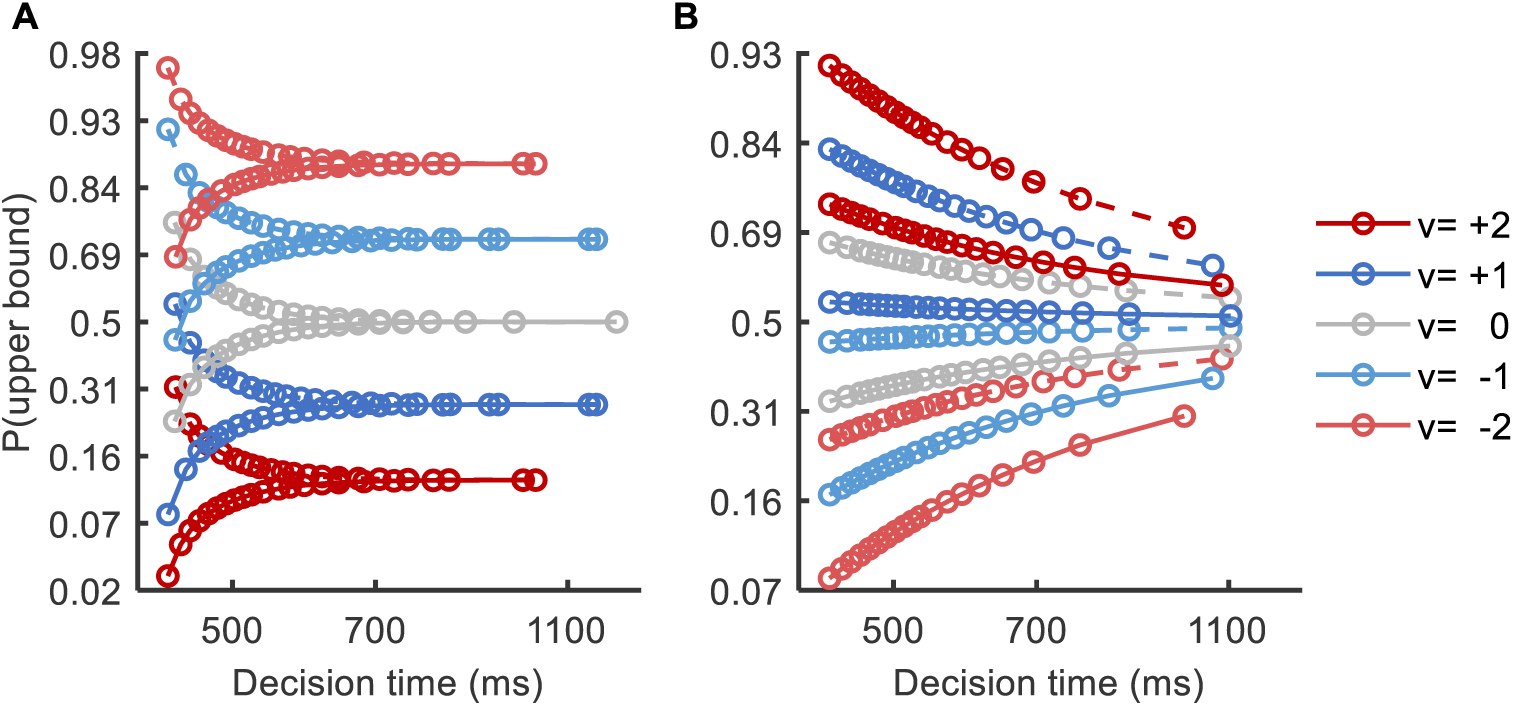
Modeling of starting point vs. variable-drift cause for reduction in bias. Shown is the percent of trials that reach the upper bound in a given decision-time bin, when the influence of adapting context (line style) is modeled using the drift diffusion model (DDM) as (**A**) a change in the starting-point (z=0.5±0.05, where the bounds are at 0 and 1), or (**B**) a change in the drift-rate (v=v±0.8) with added inter-trial variability (standard deviation of drift rate taking a value of 2). The influence of different tests orientations was modeled as different drift rates (baseline values of v, as shown in the legend). Both models show a reduction in bias as a function of time. Importantly, in the drift-rate version the context influence is obviously interchangeable with a change in the drift rate, hence there are no intersections. This illustrates the idea that when the influence of context or prior is not interchangeable with a change in evidence (e.g., in orientation of test), then this indicates that the contextual influence is time-dependent (e.g., change in starting point). Note that the exact modeling details for the contextual influence are described in the text; this figure illustrates the above idea.

## References

Clifford, C. W. G., & Rhodes, G. (2005). Fitting the mind to the world: Adaptation and after-effects in high-level vision (Vol. 2). Oxford University Press.

Dayan, P., & Daw, N. D. (2008). Decision theory, reinforcement learning, and the brain. Cognitive, Affective, & Behavioral Neuroscience, 8(4), 429–453.

Evans, J. S. B. T., & Stanovich, K. E. (2013). Dual-process theories of higher cognition: Advancing the debate. Perspectives on Psychological Science, 8(3), 223–241.

Fründ, I., Haenel, N. V., & Wichmann, F. A. (2011). Inference for psychometric functions in the presence of nonstationary behavior. Journal of Vision, 11(6).

Gold, J. I., & Shadlen, M. N. (2007). The neural basis of decision making. Annual Review of Neuroscience, 30, 535–74. https://doi.org/10.1146/annurev.neuro.29.051605.113038

Gorea, A., & Sagi, D. (2000). Failure to handle more than one internal representation in visual detection tasks. Proceedings of the National Academy of Sciences, 97(22), 12380–12384.

Green, D. M., & Swets, J. A. (1966). Signal detection theory and psychophysics.

Greenlee, M. W., Georgeson, M. A., Magnussen, S., & Harris, J. P. (1991). The time course of adaptation to spatial contrast. Vision Research, 31(2), 223–236.

Kahneman, D. (2011). Thinking, fast and slow. First edition. New York: Farrar, Straus and Giroux, 2011. Retrieved from https://search.library.wisc.edu/catalog/9910114919702121

Kaneko, S., Anstis, S., & Kuriki, I. (2017). Brief presentation enhances various simultaneous contrast effects. Journal of Vision, 17(4), 7.

Kloosterman, N. A., de Gee, J. W., Werkle-Bergner, M., Lindenberger, U., Garrett, D. D., & Fahrenfort, J. J. (2019). Humans strategically shift decision bias by flexibly adjusting sensory evidence accumulation. ELife, 8, e37321.

Magnussen, S., & Johnsen, T. (1986). Temporal aspects of spatial adaptation. A study of the tilt aftereffect. Vision Research, 26(4), 661–72.

Moran, R. (2015). Optimal decision making in heterogeneous and biased environments. Psychonomic Bulletin & Review, 22(1), 38–53.

Mulder, M. J., Wagenmakers, E.-J., Ratcliff, R., Boekel, W., & Forstmann, B. U. (2012). Bias in the brain: a diffusion model analysis of prior probability and potential payoff. Journal of Neuroscience, 32(7), 2335–2343.

Ratcliff, R. (1978). A theory of memory retrieval. Psychological Review, 85(2), 59.

Ratcliff, R., & McKoon, G. (2008). The diffusion decision model: theory and data for two-choice decision tasks. Neural Computation, 20(4), 873–922.

Ratcliff, R., & Rouder, J. N. (1998). Modeling response times for two-choice decisions. Psychological Science, 9(5), 347–356.

Ratcliff, R., Smith, P. L., Brown, S. D., & McKoon, G. (2016). Diffusion decision model: Current issues and history. Trends in Cognitive Sciences, 20(4), 260–281.

Ratcliff, R., & Tuerlinckx, F. (2002). Estimating parameters of the diffusion model: Approaches to dealing with contaminant reaction times and parameter variability. Psychonomic Bulletin & Review, 9(3), 438–481.

Schwartz, O., Hsu, A., & Dayan, P. (2007). Space and time in visual context. Nature Reviews. Neuroscience, 8(7), 522–35. https://doi.org/10.1038/nrn2155

Summerfield, C., & De Lange, F. P. (2014). Expectation in perceptual decision making: neural and computational mechanisms. Nature Reviews Neuroscience, 15(11), 745.

Tversky, A., & Kahneman, D. (1974). Judgment under uncertainty: Heuristics and biases. Science, 185(4157), 1124–1131.

Urai, A. E., De Gee, J. W., Tsetsos, K., & Donner, T. H. (2018). Choice history biases subsequent evidence accumulation. BioRxiv, 251595.

Voss, A., & Voss, J. (2007). Fast-dm: A free program for efficient diffusion model analysis. Behavior Research Methods, 39(4), 767–775.

Webster, M. A. (2011). Adaptation and visual coding. Journal of Vision, 11(5).

Wei, X.-X., & Stocker, A. A. (2017). Lawful relation between perceptual bias and discriminability. Proceedings of the National Academy of Sciences, 114(38), 10244–10249.

White, C. N., & Poldrack, R. A. (2014). Decomposing bias in different types of simple decisions. Journal of Experimental Psychology: Learning, Memory, and Cognition, 40(2), 385.

Wolfe, J. M. (1984). Short test flashes produce large tilt aftereffects. Vision Research, 24(12), 1959–64.

Zylberberg, A., Wolpert, D. M., & Shadlen, M. N. (2018). Counterfactual reasoning underlies the learning of priors in decision making. Neuron, 99(5), 1083–1097.

